# A novel iPSC model reveals selective vulnerability of neurons in Multiple Sulfatase Deficiency

**DOI:** 10.1101/2023.10.02.560535

**Authors:** Vi Pham, Livia Sertori Finoti, Margaret M. Cassidy, Jean Ann Maguire, Alyssa L. Gagne, Elisa A. Waxman, Deborah L. French, Kaitlyn King, Parith Wongkittichotee, Xinying Hong, Lars Schlotawa, Beverly L. Davidson, Rebecca C. Ahrens-Nicklas

**Author notes:** **Corresponding Author:** Rebecca C. Ahrens-Nicklas^a,b^, 3501 Civic Center Blvd, CTRB 5014, Philadelphia, PA 19104.

## Abstract

Multiple sulfatase deficiency (MSD) is an ultra-rare, inherited lysosomal storage disease caused by mutations in the gene sulfatase modifying factor 1 (*SUMF1*). MSD is characterized by the functional deficiency of all sulfatase enzymes, leading to the storage of sulfated substrates including glycosaminoglycans (GAGs), sulfolipids, and steroid sulfates. Patients with MSD experience severe neurological impairment, hearing loss, organomegaly, corneal clouding, cardiac valve disease, dysostosis multiplex, contractures, and ichthyosis. Here, we generated a novel human model of MSD by reprogramming patient peripheral blood mononuclear cells to establish an MSD iPSC line (*SUMF1* p.A279V). We also generated an isogenic control iPSC line by correcting the pathogenic variant with CRISPR/Cas9 gene editing. We successfully differentiated these iPSC lines into neural progenitor cells (NPCs) and NGN2-induced neurons (NGN2-iN) to model the neuropathology of MSD. Mature neuronal cells exhibited decreased *SUMF1* gene expression, increased lysosomal stress, impaired neurite outgrowth and maturation, reduced sulfatase activities, and GAG accumulation. Interestingly, MSD iPSCs and NPCs did not exhibit as severe of phenotypes, suggesting that as neurons differentiate and mature, they become more vulnerable to loss of *SUMF1*. In summary, we demonstrate that this human iPSC-derived neuronal model recapitulates the cellular and biochemical features of MSD. These cell models can be used as tools to further elucidate the mechanisms of MSD pathology and for the development of therapeutics.

## 1. Introduction

Multiple sulfatase deficiency (MSD) (MIM #272200) is an ultra-rare, inherited lysosomal storage disorder (LSD) characterized by the functional deficiency of all sulfatases. Sulfatases are a group of 17 enzymes that break down sulfated substrates such as glycosaminoglycans (GAGs), sulfolipids, and steroid sulfates [1]. In MSD, a severe reduction or complete absence of all sulfatase activities occurs as a result of biallelic pathogenic variants in the gene *SUMF1*, encoding the sulfatase modifying factor formylglycine-generating enzyme (FGE) [2,3]. FGE is localized to the endoplasmic reticulum and post-translationally converts a conserved cysteine residue to formylglycine in the active site of every sulfatase, a critical step for the sulfatase’s function [4]. As most sulfatases are located in the lysosomes of cells, sulfated substrates accumulate here in patients with MSD, leading to lysosomal disease manifestations.

MSD is an early-onset, progressive disease and is estimated to occur in one in 500,000 individuals worldwide [5]. Patients with MSD display complex and variable clinical presentations due to the combined deficiencies of all sulfatases. The multi-systemic nature of MSD includes symptoms from the single sulfatase deficiencies such as metachromatic leukodystrophy (MLD), several mucopolysaccharidosis subtypes (MPS II, IIIA, IIID, IVA, VI), X-linked recessive chondrodysplasia punctata type 1 (CDPX1), and X-linked ichthyosis (XLI) [5-7]. For example, MSD patients may present with psychomotor retardation and neurological deterioration, as well as vision and hearing loss, organomegaly, corneal clouding, cardiac valve disease, dysostosis multiplex, contractures, ichthyosis, and premature death [6]. Natural disease history data reveal that the mean survival of MSD patients is 13 years [7]. Currently, there are no disease-modifying therapies for MSD.

While the genetic cause of MSD has been identified and characterized, a comprehensive understanding of the underlying pathological mechanisms of MSD remains unclear. To investigate disease pathology and facilitate the development of effective treatments, appropriate models that recapitulate MSD are necessary. Animal models are often useful in elucidating disease pathophysiology and advancing novel therapeutic strategies. However, certain mouse models of LSDs may not accurately mirror the human phenotype, making them less suitable for studying disease pathology [8,9]. Multiple mouse models for MSD have been developed. The first, a *Sumf1* knock-out mouse (*Sumf1*-/-), exhibits severe growth retardation and a high mortality rate just in the first week of life. Biochemically, these *Sumf1*-/- mice lack the majority of sulfatase activities, in contrast to MSD patients who typically retain residual activity [10]. Subsequently, two less-severe mouse models were developed by introducing common pathogenic variants of MSD into the mouse *Sumf1* gene (p.S153P or p.A277V). While these novel mouse models recapitulate MSD biochemically, they do not necessarily demonstrate all the severe neurobehavioral phenotypes observed in patients [11]. In addition, the regulation of gene expression can vary across species. For example, a key microRNA (miR-95) that is shown to regulate *SUMF1* expression is not present in mice [12]. Therefore, there is an unmet need for MSD models that reproduce human phenotypes.

Recent technological advances in generating induced pluripotent stem cells (iPSCs) and isogenic controls have enabled the establishment of iPSC-based disease models, including for LSDs. iPSCs are ideal cellular models for studying human disease because they are reprogrammed from patient somatic cells, thereby harboring natural disease-causing variants within the patient’s genetic background. Moreover, CRISPR/Cas9 gene editing can be used to precisely modify parental iPSC lines, allowing for the production of corrected isogenic control iPSC lines for appropriate comparison [13]. Additionally, iPSCs not only exhibit sustained growth in vivo, but can also be differentiated into various somatic cells, including blood cells, hepatocytes, cardiomyocytes, and pancreatic beta cells [14-17]. Importantly, iPSCs can be differentiated into cellular lineages that are difficult to obtain from patients, such as neural progenitor cells (NPCs) or neurons [18]. Therefore, iPSCs play an important role in investigating neurological phenotypes and neurodegeneration, which are central to many LSDs, including MSD.

Neurodegeneration is a prominent feature of most LSDs, including MSD [19]. Patients with LSDs experience neurological manifestations, such as developmental delay, seizures, or psychiatric problems. Specifically for MSD, patients present with neurologic symptoms as the most common first clinical feature and neurological disability is universal [6,7]. These phenotypes suggest that cells of the central nervous system (CNS) are particularly sensitive to dysfunctional cellular clearance in MSD. However, the specific impact of lysosomal storage on neuronal function and degeneration remains unclear. It is imperative to uncover the mechanisms of neuronal dysfunction in MSD to develop efficient therapies that address the neurological symptoms of the disease.

Here, we generate a novel human iPSC model of MSD that can be used to examine the effects of the pathological storage on neurodevelopment, survival, and function of neural cells. To do so, we first reprogrammed peripheral blood mononuclear cells from an MSD patient (homozygous for the *SUMF1* p.A279V) into iPSCs. Next, we generated an isogenic control iPSC line by specifically correcting the variant with gene editing. Using distinct protocols, we successfully differentiated these iPSCs into two neural models: 1) neural progenitor cells and 2) early neurons. We used these neuronal models to explore MSD neural pathology and found that both models recapitulated MSD cellular and biochemical disease phenotypes, including sulfatase deficiencies and GAG accumulation. Notably, these phenotypes became more severe with neuronal maturation. Overall, we show that MSD iPSC-derived NPCs and neurons are valuable cell-based models to assess disease neuropathology and may be used to develop potential treatment strategies targeting the neurological symptoms of MSD. Furthermore, this novel human-derived MSD iPSC model will be an important tool for studying the mechanisms underlying this multi-systemic disease across various cell types.

## 2. Materials and Methods

### 2.1. Generation of *SUMF1* patient A279V iPSC line

The MSD iPSC line was generated from an MSD patient (homozygous for *SUMF1* p.A279V) following a previously published protocol [20]. Briefly, cellular reprogramming was performed using ficoll-purified mononuclear cells from whole blood that were expanded to erythroblasts for transduction with Sendai viral vectors expressing human OCT3/4, SOX2, KLF4, and cMYC according to manufacturer’s instructions (ThermoFisher Scientific). Transduced cells were plated on culture dishes containing irradiated murine embryonic fibroblasts (MEFs) and maintained in human embryonic stem cell (HES) medium consisting of DMEM/F12 (50:50; Mediatech) supplemented with 20% knock-out serum replacement (ThermoFisher Scientific), 10−4 M mercaptoethanol (Sigma), and 10 ng/mL bFGF (R&D Systems). The medium was replenished every 2-3 days for 3 weeks. Cells were maintained in these conditions until uniform colonies were generated and iPSC colonies were mechanically isolated for expansion on MEFs. Cells were subsequently transitioned to feeder-free cultures and maintained with mTeSR1 (StemCell Technologies) on hESC qualified Matrigel (Corning). Feeder-free iPSC stocks were cryopreserved in 90% FBS/ 10% DMSO at a minimum of 2 passages after feeder-free transition. Feeder-free stocks were passaged in mTeSR1 prior to the initiation of differentiation. The authentication of each clone confirming identity to the original patient cells was performed by DNA fingerprinting using PCR. Mutation verification was also performed on genomic DNA by PCR amplification and sequence analyses performed. Karyotype analysis was outsourced to Cell Line Genetics (Madison, WI). Stemness surface markers were performed by flow cytometry and mycoplasma was tested by PCR.

### 2.2. Generation of *SUMF1* corrected A279V iPSC line

The isogenic control iPSC line was obtained by correcting the pathogenic variant c.836C>T in the *SUMF1* patient A279V iPSC line using a CRISPR/Cas9 system previously-described [13]. In brief, the p.A279V homozygous mutation was corrected by homology directed repair using one 100 bp single-stranded oligonucleotide (ssODN). Silent mutations were included in the guide RNA (gRNA) sequence to prevent recutting, and a unique restriction site was introduced to aid in screening targeted clones. Three gRNAs were designed within ±20bp of the editing site and tested for efficiency. The gRNA with the sequence 5’-ACTGGTGAGGATGGCTTCCAAGG-3’ was ultimately used to target the 279 residue of *SUMF1*. The ssODN was transfected along with CAS9-GFP and the gRNA. The GFP+ cells were sorted and plated at clonal density. Single colonies were screened using PCR followed by restriction digest. Sequencing was performed to confirm editing.

### 2.3. Differentiation of iPSCs into neural progenitor cells

Differentiation of iPSCs into NPCs was initiated, as previously described [21] with indicated modifications. Cultures were treated with daily media changes containing SB431542 (10 μM; Tocris), LDN193189 (1 μM, Tocris), and endo-IWR1 (1.5 μM; Tocris) and supplemented with B27 without vitamin A (Invitrogen) and passaged at days 4 and 8 of differentiation. After 8 days of daily treatment, forebrain neural ectoderm was confirmed by expression of FOXG1, SOX2, and PAX6 by flow cytometry. From days 8 to 14 of differentiation, NPCs were expanded in Invitrogen neural expansion media, containing Neural Induction Supplement in Advanced DMEM/F12 and Neurobasal Medium (Invitrogen, per manufacturer instructions). On Day 14, NPC identity was confirmed by expression of Forse-1 by flow cytometry and by rosette morphology. Day 14 NPCs were cryopreserved in 90% FBS/ 10% DMSO for future use. Brightfield images were captured with an EVOS xl, version 1.0.198. iPSC images were captured with a 10X objective, and NPC images were captured with a 20X objective.

### 2.4. Differentiation of iPSCs into neurons

NGN2-iN lentiviruses were produced by polyethylenimine (Polysciences, PEI, 23966-2,) in HEK 293T cells (ATCC, VA) using 3rd generation lentiviral packaging plasmids pMDLg/pRRE (Addgene Inc., Plasmid 12251) and pRSV-RV (Addgene Inc., Plasmid 12253), the VSC-G envelope expressing plasmid (Addgene Inc., Plasmid 12259), in addition to the plasmid pLVX-UbC-rtTA-Ngn2:2A:EGFP (Addgene Inc., Plasmid 127288). Plasmids were removed after 8 hours, and HEK 293T cells were replenished with mTeSR™ Plus medium (StemCell Technologies). After 48 hours, conditioned media was collected and passed through a 0.45 µm filter to remove cell debris.

iPSCs were differentiated into NGN2-iN by following established protocols [22,23]. Briefly, MSD3 iPSCs were transduced with lentiviral constructs in mTeSR Plus medium containing 10 µM Y-27632 (ROCK inhibitor; Tocris, 1254) for 24 hours. The transduced cells were selected with 0.5-1 µg/mL puromycin, and individual colonies were selected by PCR verification for the presence of the puromycin resistance gene. Selected cells were differentiated using doxycycline-induced expression of NGN2. iPSC cells were dissociated using Accutase (Sigma-Aldrich, A6964) and resuspended in pre-differentiation Medium (Knockout DMEM/F12 (Gibco, 12660-012), N2 supplement (Gibco, A1370701), MEM Non-Essential Amino Acids (Gibco, 11140-050)) supplemented with 10 µM ROCK inhibitor, 0.2 µg/mL Mouse Laminin (ThermoFisher Scientific, 23017-015), 10 ng/mL NT-3 (PreproTech, 450-03), 10 ng/mL BDNF (PreproTech, 450-02), and 2 μg/mL doxycycline hydrochloride (Sigma-Aldrich, D3072) and plated in hESC-Qualified, LDEV-Free Matrigel-Matrix (Corning, 354277) coated plates to induce expression of NGN2. Pre-differentiation media was supplemented with laminin, NT-3, BNDF, and doxycyline hydrochloride and replaced each day. ROCK inhibitor was added to the media for the first day after replating. After 3 days, pre-differentiated cells were dissociated as detailed and replated in Neuronal Medium (Neurobasal Plus Medium (Gibco, A3582901), N2, B27 (Gibco, 17504-044), MEM Non-Essential Amino Acids, Glutamax supplement (Gibco, 35050-061)) supplemented with 2 μg/mL doxycycline hydrochloride, 10 ng/mL NT-3, 10 ng/mL, 1 µg/mL Mouse Laminin on plates coated with 100 µg/mL poly-L-ornithine (Sigma-Aldrich, P3655) and Matrigel. On day 7, half of the media was exchanged with fresh Neuronal Medium supplemented without doxycycline.

### 2.5. Flow cytometry

Cells were retrieved in Accutase, diluted with DPBS, and then centrifuged at 3000 x g, followed by resuspension into 1.6% paraformaldehyde and incubated for 30 min at 37°C with agitation. Cells were subsequently washed once with DPBS and resuspended into FACS Buffer (DPBS containing 0.5% BSA (Jackson ImmunoResearch) and 0.05% sodium azide) and stored at 4°C until flow cytometry analysis. Flow cytometry analysis was performed with permeabilization of cells with saponin buffer (diluted to 1X in water; Biolegend cat# 421022) and 1 hr incubation of the following antibodies diluted in saponin buffer: SOX2 (1:300, Cell Signaling technology, 3569), PAX6 conjugated to Alexa 647 (1:20, BD, 562249), FOXG1 (1:300, Abcam, ab196868), or maintained in FACS buffer: Forse-1 (1:100, DSHB), IgM isotype control (1:100, Santa Cruz, sc-3881). Cells were subsequently washed with saponin buffer or FACS buffer and incubated for 1 hr with secondary antibodies: goat anti-rabbit Alexa 488 (1:500, Jackson Immunoresearch, 111-545-144) or goat anti-mouse IgM-conjugated Alexa 488 (1:500, ThermoFisher, A21042). Cells were subsequently washed and resuspended into FACS buffer. Cells were analyzed using a CytoFLEX flow cytometer (Beckman Coulter) and the FlowJo software program (BD).

### 2.6. Immunocytochemistry

NGN2-induced neurons were grown on poly-L-ornithine and Matrigel coated 6-well plates. At the time of collection, cells were gently washed twice with PBS and fixed at room temperature for 20 minutes with 4% paraformaldehyde. Wells were washed three times with PBS and left at 4°C until immunocytochemistry (ICC) was performed (up to 1 week). Wells were permeabilized for 10 minutes with 0.1% Triton X-100 (Sigma-Aldrich, T8787) in PBS and blocked for 2 hours at room temperature with 5% Normal Goat Serum (Jackson Immuno, 005-000-121) in PBS. After blocking, primary antibodies (Rabbit anti-MAP2, Millipore, AB5622, 1:800; Mouse anti-Tuj1, R&D Systems, mab1195, 1:500) were diluted in PBS and added to the wells to incubate overnight at 4°C while gently rocking. After primary incubation, wells were briefly washed with PBS and secondary antibodies (Donkey Anti-Rabbit 555, Invitrogen, A32794, 1:1000; Donkey Anti-Mouse 647, Invitrogen, A32787, 1:1000) were diluted in PBS and added to wells to incubate for 1 hour at room temperature while covered from light. DAPI (ThermoFisher, D1306) was then diluted 1:10,000 in PBS and added to the wells for 10 minutes at room temperature. Finally, wells were washed and covered with PBS. Imaging was performed with a Keyence BZ-X800.

### 2.7. Western blotting

Total protein concentrations were measured in cell lysates using the Pierce BCA Protein Assay Kit (Thermo Fisher Scientific). A total of 15 µg of total protein fraction for each sample was separated by electrophoresis using NuPAGE™ 4-12% Bis-Tris Protein Gel (Thermo Fisher Scientific). Proteins were transferred to a polyvinylidene difluoride membrane using an XCell SureLock Mini-Cell Electrophoresis System (Thermo Fisher Scientific). The membrane was blocked using 5% milk for 45 minutes at room temperature. After briefly rinsing with Tris-Buffered Saline with Tween-20, the membrane was incubated overnight at 4°C with one of the following antibodies: FGE (R&D Systems, MAB2779, 1:1000), LAMP1 (DSHB, H4A3, 1:115), or Vinculin (Abcam, ab91459, 1:2000). The membrane was then washed with Tris-Buffered Saline with Tween-20 and incubated in secondary antibody (Goat Anti-Rabbit IgG H&L (HRP), Abcam, ab6721; Goat Anti-Mouse IgG H&L (HRP), Abcam, ab205719) for 1.5 hours at room temperature. Clarity Max™ Western ECL Substrate (BioRad 1705062) and ChemiDoc XRS+ Imaging System (BioRad) were used for visualization.

### 2.8. RNA isolation, cDNA synthesis, and qPCR

Total RNA was isolated from cells using Qiagen RNeasy Mini kit (Qiagen, 74104) according to manufacturer’s instructions. Quantification and quality of isolated RNA was assessed by Nanodrop measurements. cDNA was prepared using the High Capacity RNA-to-cDNA kit (Applied Biosystems, 4387406). The TaqMan gene expression assays performed for target genes *SUMF1* (ID: Hs00399783_m1) and *NEUROD1* (ID: Hs01922995_s1) were normalized to the average expression of *RPLP0* (ID: Hs00420895_gH), *RNA18S5* (ID: Hs03928989_g1), and *GAPDH* (ID: Hs02786624_g1) as housekeeping genes. qPCR was conducted using TaqMan™ Fast Advanced Master Mix (Applied Biosystems, 4444556) on a Quantstudio 5 Real-Time PCR System (Applied Biosystems).

### 2.9. Cell proliferation analysis

For cell proliferation assays, 500,000 cells were plated into one-well (in triplicate) of an Geltrex-coated 6-well plate on Day 0. One and three days after plating, cells were counted using an automated Countess II cell counter (Thermo Fisher).

### 2.10. Sulfatase activity assays

ARSA and ARSB activities were determined from whole cell lysates following previously published protocols [24]. Briefly, 50 µg of protein extract was incubated at 37°C for 1 hr with the artificial p-nitrocatecholsulfate substrate (Santa Cruz Biotechnology, sc-238927A) and either sodium pyrophosphate (ARSA) or barium acetate (ARSB). The absorbance was measured at 515 nm in a BioTek Epoch Microplate Spectrophotometer (Agilent). ARSA and ARSB units are defined as the amount of 4-nitrocatechol liberated in 1 hr per mg of total protein. SGSH activity was measured from whole cell lysates following a two-step protocol [25] and using 4MU-αGlcNS (Chem-Impex Int’l Inc, Cat #30694) as fluorogenic substrate and with 4-methylumbelliferone (Sigma Aldrich) as standard. A unit is defined as the amount that will liberate 1 nmol 4-MU at 37°C from the respective fluorogenic substrate in 17 hr.

### 2.11. Glycosaminoglycan quantification

GAG fragments (i.e., sulfated oligosaccharides) in cells were measured using a modified protocol as described [26,27]. Chondroitin disaccharide di-4S (dUA-GalNAc-4S, Biosynth) was used as the internal standard (IS). Cell lysates equivalent to 100 µg protein were used. The UPLC-MS/MS analysis was carried out on a Waters TQ-S tripe quadrupole mass spectrometry coupled to an ACUIQTY Classic UPLC system. Three previously reported markers, HexN-UA-1S (MPS-IIIA), GlcNAc-6S (MPS-IIID), and GalNAc-4S (MPS-VI), were analyzed [26,27]. Two novel markers, UA-HexN-UA-2S (509.8 -> 422.7) and HexN-UA-HexNAc-UA-2S (611.4 -> 524.1), associated with MPS-I/II and MPS-IIIA, respectively, were also included. The apparent concentration was calculated by multiplying the response ratio of the analyte and the IS by the amount of IS added (fmol), then dividing by the amount of protein (µg).

### 2.12. Image quantification

Western Blots and immunofluorescence images were quantitatively analyzed using Fiji software [28]. Statistical analysis was performed using Prism (GraphPad software, San Diego, USA).

### 2.13. Statistical analyses

Results were calculated from three independent biological replicates (except NPC data which had one biological replicate), and replicate measurements were summarized as mean values in respective calculations. All statistical analysis was performed using Prism (GraphPad software, San Diego, USA). Comparison of two independent groups was done using an unpaired *t*-test. Two-way ANOVA followed by Bonferroni’s multiple comparison test was used to compare groups (isogenic control vs. MSD and differentiation cell type). Data were expressed and displayed as mean and standard error. Significance levels were displayed as follows: *p < 0.05, **p > 0.01, ***p < 0.001, ****p < 0.0001 for differences indicated in the figure legends. Differences were not statistically significant if not indicated.

## 3. Results

### 3.1. Generation of patient-derived iPSCs and gene editing to generate isogenic controls

To generate an iPSC model of MSD, peripheral blood mononuclear cells were isolated from the whole blood of an MSD patient that was homozygous for the recurrent pathogenic variant *SUMF1* c.836C>T, p.A279V. Cells were expanded to erythroblasts before being reprogrammed using non-integrating Sendai viral vectors expressing the human factors OCT3/4, SOX2, KLF4, and cMYC as described [23] (Fig. 1A). The mutation (*SUMF1* c.836C>T) was confirmed by Sanger sequencing on genomic DNA and the established MSD iPSC line was karyotypically normal (Figs. 1B, S1A). Mycoplasma testing was negative and continued to be negative throughout the course of the experiment (Fig. S1B). We found that MSD iPSCs possessed embryonic stem cell-like morphology and the expression of typical stemness markers (SSEA3, SSEA4, TRA160, and TRA181) (Fig. S2A).

**Figure 1.**
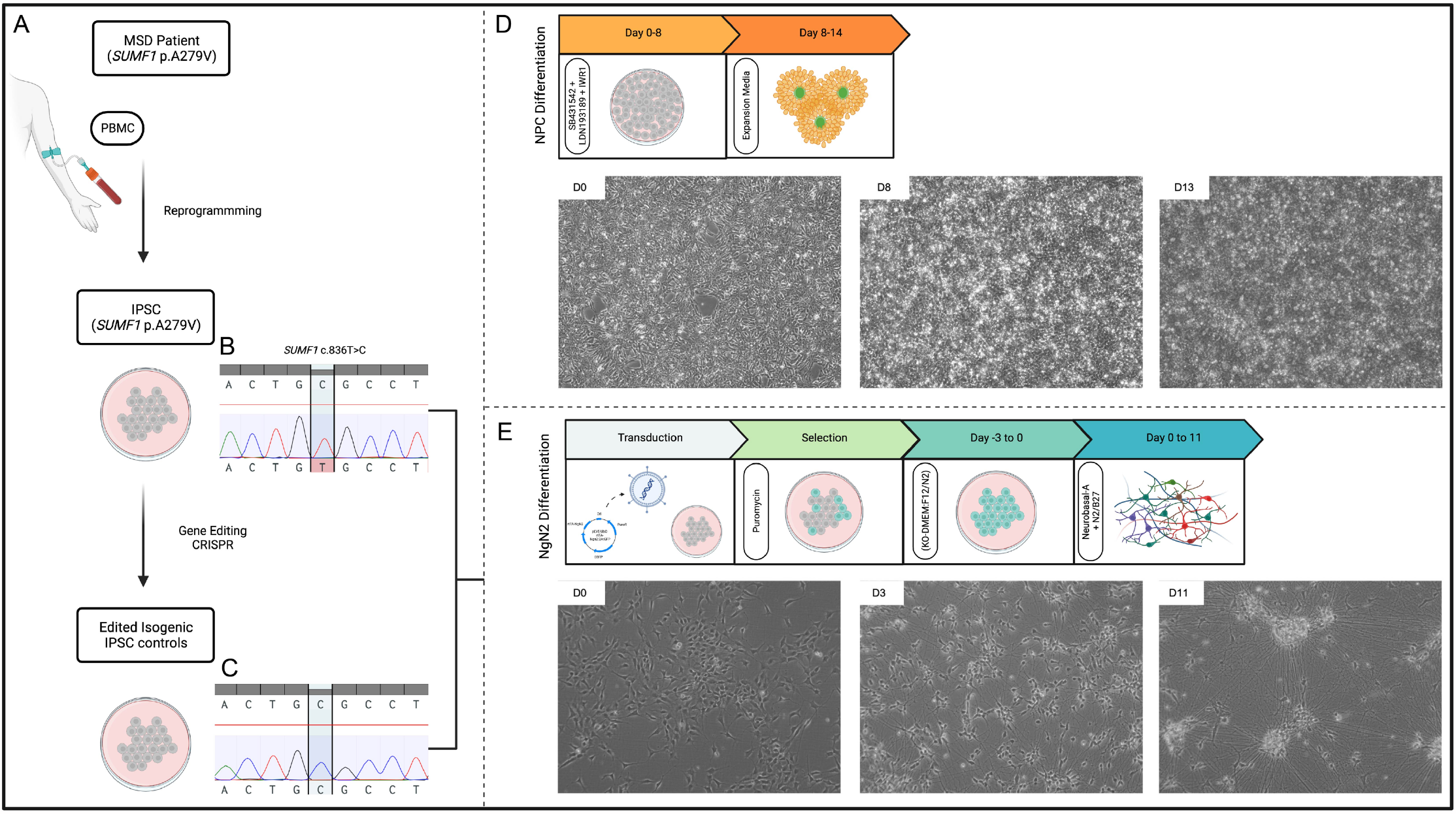
MSD iPSC generation and differentiation into neuronal models. **A**. Schematic of iPSC reprogramming from MSD patient peripheral blood mononuclear cells (PBMCs) and generation of isogenic control iPSC line. **B**. Chromatogram confirming patient variant *SUMF1* c.836C>T in MSD iPSC line after reprogramming. **C**. Chromatogram confirming homozygous correction of patient variant *SUMF1* c.836T>C in isogenic control iPSC line after gene editing with CRISPR/Cas9. **D**. NPC differentiation timeline with media and supplements used throughout the protocol indicated. Representative brightfield images confirming characteristic NPC morphology at Days 0, 8, and 13 (D0, D8, and D13) of the differentiation protocol. **E**. NGN2-iN differentiation timeline with media and supplements used throughout the protocol indicated. Representative brightfield images confirming characteristic neuronal morphology at Days 0, 3, and 11 (D0, D3, and D11) of the differentiation protocol. Created with BioRender.com.

We next generated an isogenic control iPSC line in order to reduce variation in the genetic background between groups. We used CRISPR/Cas9 to correct the pathogenic c.836C>T in the MSD iPSC parental line. The correction was validated by Sanger sequencing, CNV analysis of this iPSC line showed a normal karyotype, and mycoplasma testing was negative (Figs. 1C, S1C,D). The corrected isogenic control iPSCs also expressed the stemness markers SSEA3, SSEA4, TRA160, and TRA181 as detected by flow cytometry (Fig. S2B).

### 3.2. Differentiation of iPSC to NPCs and NGN2-induced neurons (NGN2-iNs)

We hypothesized that neurons are a particularly vulnerable cell population in MSD due to the severity of the neurological phenotypes experienced by patients. In order to examine the neuropathology of MSD patients, we successfully differentiated MSD and isogenic control iPSCs into NPCs and neurogenin-2-induced neurons (NGN2-iNs) (Figs. 1D,E). These neural cell types were selected as models because NPCs represent early CNS cells that can give rise to both glial and neuronal cell types, whereas NGN2-iNs represent more mature neurons and may bypass NPC-like stages [22].

To produce the NPC models, we differentiated MSD and isogenic control iPSCs using a 14-day protocol as previously described [21]. Flow cytometry analysis revealed that cells expressed the classical NPC markers SOX2, PAX6, and FOXG1 at Day 8 of differentiation. By Day 14, cells expressed Forse-1 and rosette morphology confirming their NPC phenotype (Fig. S2, 1D). We found that MSD NPCs demonstrated normal differentiation and rosette morphology, as shown by brightfield imaging (Fig. 1D).

We additionally generated a more mature neuronal model of MSD through direct NGN2-mediated differentiation of iPSCs. Neurons derived from iPSCs by overexpressing NGN2 have been extensively used to investigate the processes of neuronal differentiation and to create models of neurological disorders [29]. We produced MSD and isogenic control NGN2-iNs by differentiating iPSCs using established protocols [26,27]. NGN2-iNs were collected for analyses at Days 3 (D3 NGN2-iN) and 11 (D11 NGN2-iN) of differentiation in order to compare phenotypes during neuronal development. iPSCs demonstrated neurite outgrowth by Day 3 and had elaborate processes by Day 11 (Fig. 1E). These findings indicate that MSD iPSCs can successfully differentiate into NPCs and NGN2-iNs to produce neural models.

### 3.3. *SUMF1* transcript levels and FGE protein expression increase throughout neuronal differentiation in isogenic controls but not MSD cells

The majority of pathogenic *SUMF1* variants are missense variants that ultimately cause misfolding of the FGE protein [6,30]. Improper folding leads to the early degradation of these FGE variants when misfolded FGE interacts with protein disulfide isomerase in the endoplasmic reticulum [31]. To determine if our MSD iPSCs and differentiated neural cells exhibited this characteristic molecular feature of the disease, we measured FGE protein expression by Western blot at different stages of differentiation: iPSCs, NPCs, D3 NGN2-iNs, and D11 NGN2-iNs (Figs. 2A, S3A). Interestingly, we found that FGE protein expression did not significantly differ between MSD cells and isogenic controls at iPSC, NPC, and D3 NGN2-iN stages (Figs. 2B, S3A). However, as iPSC-derived neurons matured, differences in FGE protein expression became more apparent. D11 MSD NGN2-iNs showed a trend towards decreased FGE expression as compared to their isogenic controls, although not statistically significant. These findings suggest that there may be a preferential degradation of variant FGE in specific cell types, such as neurons.

**Figure 2.**
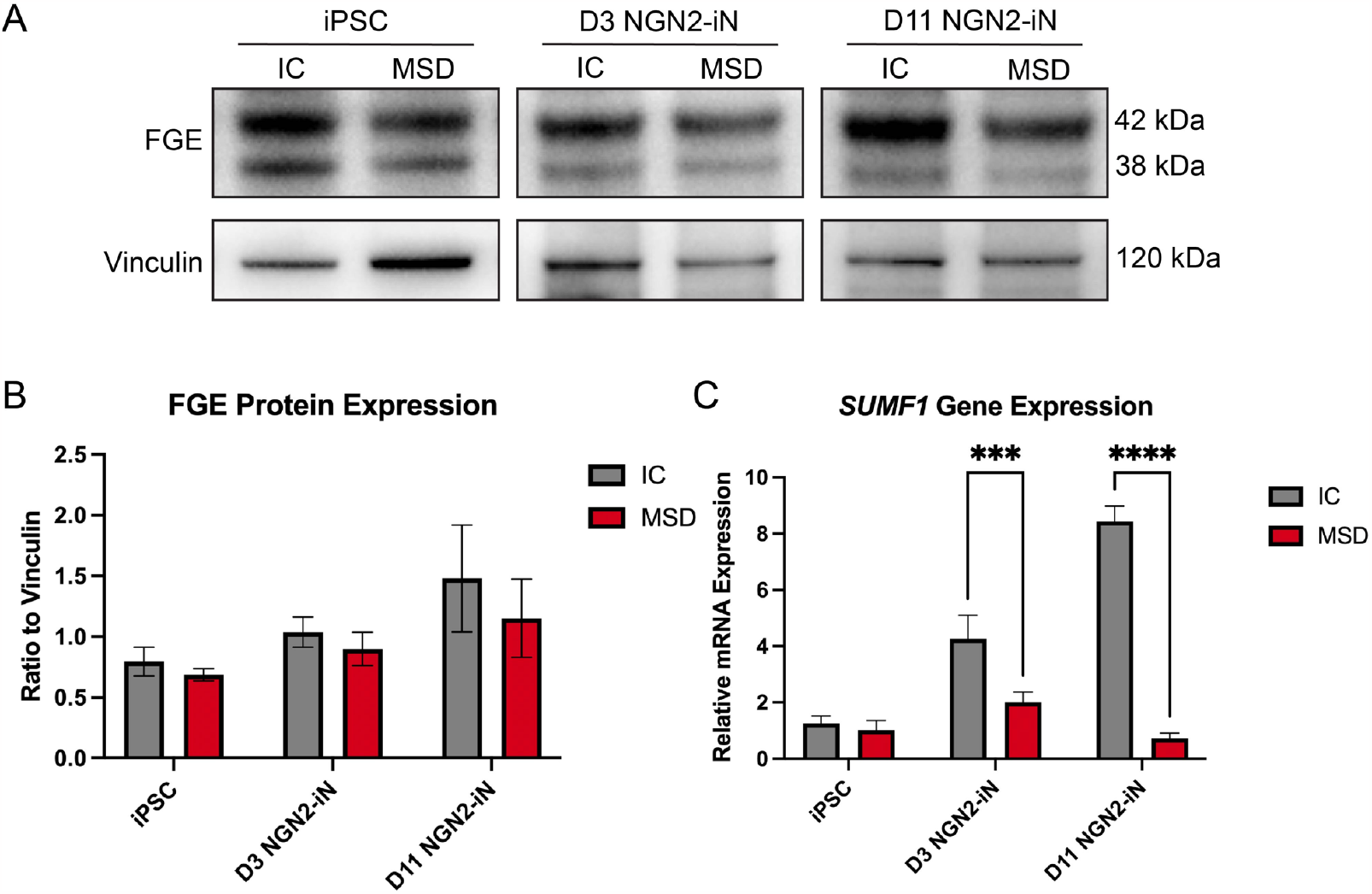
FGE protein expression and *SUMF1* gene expression was reduced in MSD NGN2-iNs. **A**. Western blot analysis of FGE protein expression in MSD and isogenic control (IC) cells at different stages of neuronal maturation: iPSCs, Day 3 NGN2-iNs (D3 NGN2-iN), and Day 11 NGN2-iNs (D11 NGN2-iN). 42 kDa band represents the glycosylated form of FGE and 38 kDa band represents the non-glycosylated form of FGE. Vinculin was used as a loading control. **B**. Quantification of FGE Western blot analysis. FGE protein expression was quantified as a ratio to the vinculin loading control per sample. Data are mean +/-s.e.m. from 3 biological replicates. **C**. Quantification of *SUMF1* gene expression in MSD vs. isogenic control iPSCs, Day 3 NGN2-iNs (D3 NGN2-iN), and Day 11 NGN2-iNs (D11 NGN2-iN). Relative *SUMF1* mRNA expression was normalized to the average expression of three different housekeeping genes as controls (*GAPDH, RNA18SN5*, and *RPLP0*). Data are mean +/-s.e.m. from 3 biological replicates. ***p < 0.001 and ****p < 0.0001.

To compare the differences in *SUMF1* gene expression, we performed qPCR on MSD iPSCs, NPCs, D3 NGN2-iNs, and D11 NGN2-iNs (Figs. 2C, S3B). Interestingly, we found an upregulation of *SUMF1* gene expression throughout differentiation in isogenic control iPSCs, reaching the highest levels in Day 11 NGN2-iNs. Conversely, *SUMF1* levels remained constant during the neuronal differentiation of MSD iPSCs. At the iPSC and NPC stages of differentiation, *SUMF1* transcript levels did not differ between MSD cells and isogenic controls. However, as neuronal differentiation progressed, MSD NGN2-iNs exhibited significantly less *SUMF1* expression when compared to their isogenic controls, both at Days 3 and 11 of differentiation. Taken together, these results indicate that *SUMF1* transcript and FGE protein levels increase with neuronal maturation and that these levels are significantly reduced in MSD NGN2-iNs.

### 3.4. iPSCs, NPCs, and NGN-iNs from MSD patients display growth abnormalities, lysosomal stress, and maturation defects

To characterize disease phenotypes in these cellular models of MSD, we performed proliferation assays on MSD iPSCs (Fig. 3A,B). We measured the growth rate of MSD iPSCs as compared to isogenic control iPSCs by comparing the number of cells at various time points after initial plating. MSD proliferated more slowly as compared to that of isogenic control iPSCs. At Day 0, we plated equal numbers of cells for both groups. One day after plating (Day 1), there was a 2.3-fold greater number of isogenic control iPSCs as compared to MSD iPSCs (Fig. 3B). The reduced number of MSD iPSCs at Day 1 suggests either cell death following replating or a failure of the MSD cells to attach. However, the difference in fold increase persists at Day 3, as there was about a 1.3-fold greater number of isogenic control iPSCs than MSD iPSCs in the well (Fig. 3B). These findings indicate a reduced proliferative capability of MSD iPSCs as compared to isogenic controls.

**Figure 3.**
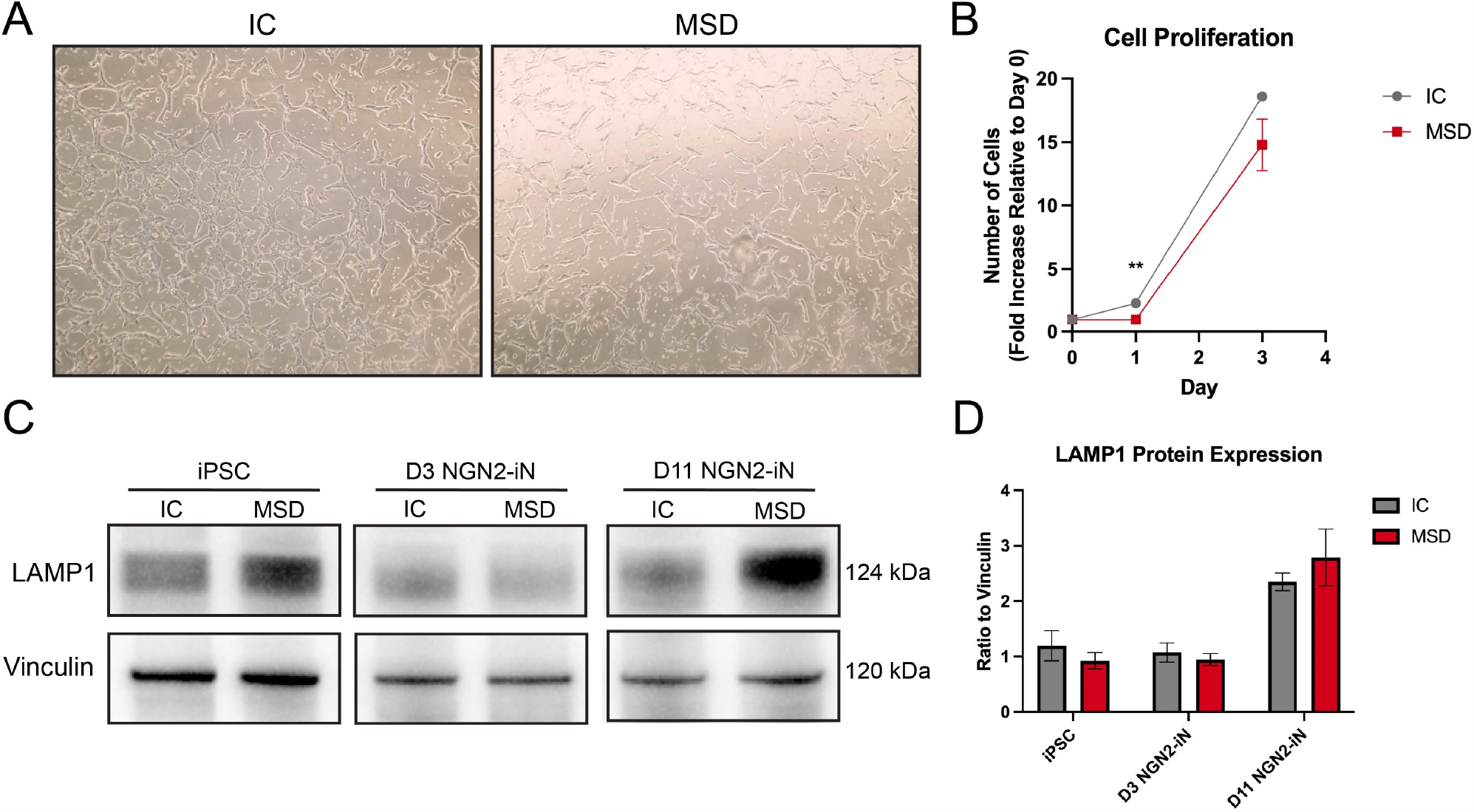
MSD cells displayed decreased rates of proliferation and increased lysosomal stress. **A**. Representative brightfield images of confluency for isogenic control (IC) and MSD iPSCs one day after initial plating. **B**. Quantification of cell growth over time for isogenic control and MSD iPSCs. The number of cells is reported as the fold increase relative to the number of cells at Day 0. **C**. Western blot analysis of LAMP1 protein expression in MSD and isogenic control (IC) cells at different stages of neuronal maturation: iPSCs, Day 3 NGN2-iNs (D3 NGN2-iN), and Day 11 NGN2-iNs (D11 NGN2-iN). Vinculin was used as a loading control. **D**. Quantification of LAMP1 Western blot analysis. LAMP1 protein expression was quantified as a ratio to the vinculin loading control per sample. Data are mean +/-s.e.m. from 3 biological replicates. **p < 0.01.

Enlarged lysosomes are a hallmark feature of cellular pathophysiology in lysosomal disorders including MSD [32]. We next assessed the lysosomal phenotypes of MSD iPSCs, NPCs, and NGN2-iN cell models by measuring the expression of lysosomal-associated membrane protein 1 (LAMP1) by Western blot (Figs. 3C, S3C). Consistent with FGE protein expression, MSD cells did not exhibit significant differences in LAMP1 expression at the iPSC, NPC, and Day 3 NGN2-iN stages of differentiation (Figs. 3D, S3C). However, at Day 11 of differentiation, MSD NGN2-iNs showed a trend towards increased LAMP1 expression as compared to isogenic controls, although not significant. These findings suggest that pushing the neurons further into the differentiation process may increase lysosomal stress on the cells, indicative of a selective vulnerability of neurons to MSD pathology.

We next sought to characterize phenotypic differences in neuronal growth and maturation of MSD cells by evaluating the levels of two neuronal markers, MAP2 and Tuj1, in MSD NGN2-iNs (Fig. 4). MAP2 belongs to the microtubule-associated protein family and is involved in microtubule assembly, an essential step in neurogenesis. This neuron-specific cytoskeletal protein is enriched in dendrites and enables the characterization of early neuronal development. Tuj1 is a class III member of the beta tubulin protein family and is also found almost exclusively in neurons, making it a classical marker of mature neurons. We performed ICC for MAP2 and Tuj1 in MSD NGN2-iNs at Days 3 and 11 of differentiation and compared it to that of isogenic control NGN2-iNs. Both MSD and isogenic control NGN2-iNs express MAP2 and Tuj1 confirming their successful differentiation into neurons (Fig. 4A). Notably, MSD NGN2-iNs displayed differences in MAP2 and Tuj1 expression levels as compared to isogenic controls (Figs. 4B,C). At Day 3 of NGN2-iN differentiation, MSD cells showed a trend towards decreased MAP2 and Tuj1 expression compared to isogenic controls suggesting delayed neurite outgrowth. These differences increased in severity as differentiation progressed, with Day 11 MSD NGN2-iNs exhibiting significantly reduced levels of both markers compared to isogenic controls. Additionally, as seen in the immunofluorescence imaging, Day 11 MSD NGN2-iNs show impairments in neuronal organization and structural formation as compared to isogenic control NGN2-iNs (Fig. 4A, third and fourth columns). Isogenic control iPSCs differentiated into NGN2-iNs that separated into distinct clusters of cell bodies with protruding neurites in a radial organization. On the other hand, MSD iPSCs differentiated into NGN2-iNs with a less-structured morphology (fewer distinct clusters and disorganized neural processes). In summary, MSD neurons have impaired neurite outgrowth at early stages of differentiation and abnormal neurite organization throughout maturation.

**Figure 4.**
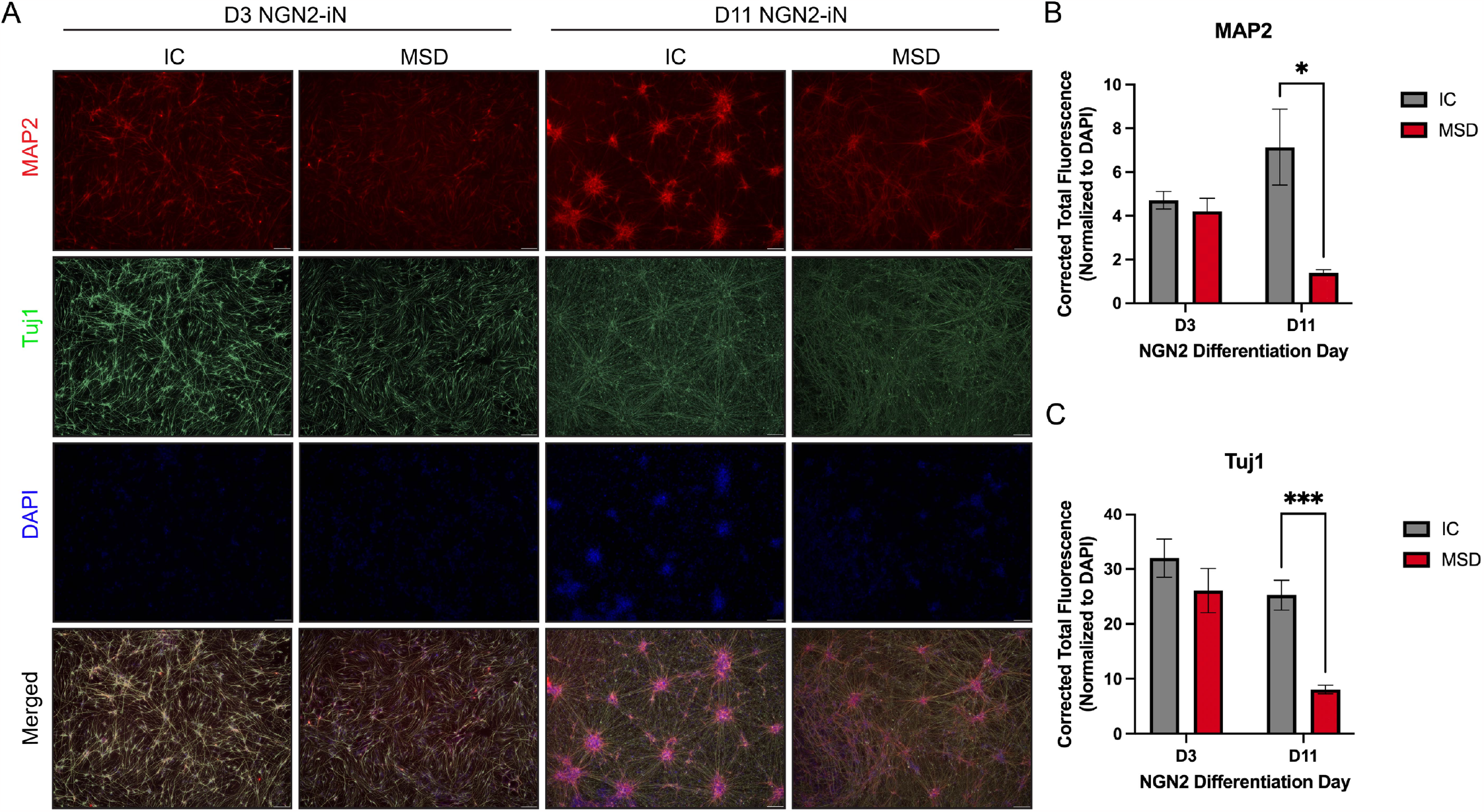
MSD NGN2-iNs exhibited impaired neurite outgrowth and maturation. **A**. Representative immunofluorescence images of MAP2 and Tuj1 staining of isogenic control (IC) and MSD Day 3 and Day 11 NGN2-iNs (D3 NGN2-iN, D11 NGN2-iN). Labeling with anti-MAP2 antibody (red fluorescence), anti-Tuj1 antibody (green pseudo-fluorescence), and DAPI (nuclei, blue). Scale bar = 100 µm. **B**. Quantification of MAP2 immunofluorescence staining normalized to DAPI. **C**. Quantification of Tuj1 immunofluorescence staining normalized to DAPI. Data are mean +/-s.e.m. (10 regions of interest x 2-3 biological replicates). *p < 0.05 and ***p < 0.001.

### 3.5. Sulfatase activities are deficient in MSD neuronal cells

These cellular phenotypes prompted us to elucidate the biochemical differences underlying the cellular phenotypes exhibited by MSD iPSCs, NPCs, and NGN2-iNs. FGE activates all sulfatases in the cell. Therefore, pathogenic variants in *SUMF1*, encoding FGE, lead to the functional deficiency of all sulfatases in patients with MSD. To characterize how the pathogenic FGE variant influences specific sulfatase activities, we measured the activities of three downstream sulfatases: arylsulfatase A (ARSA), arylsulfatase B (ARSB) and sulfamidase (SGSH) (Figs. 5, S3D).

**Figure 5.**
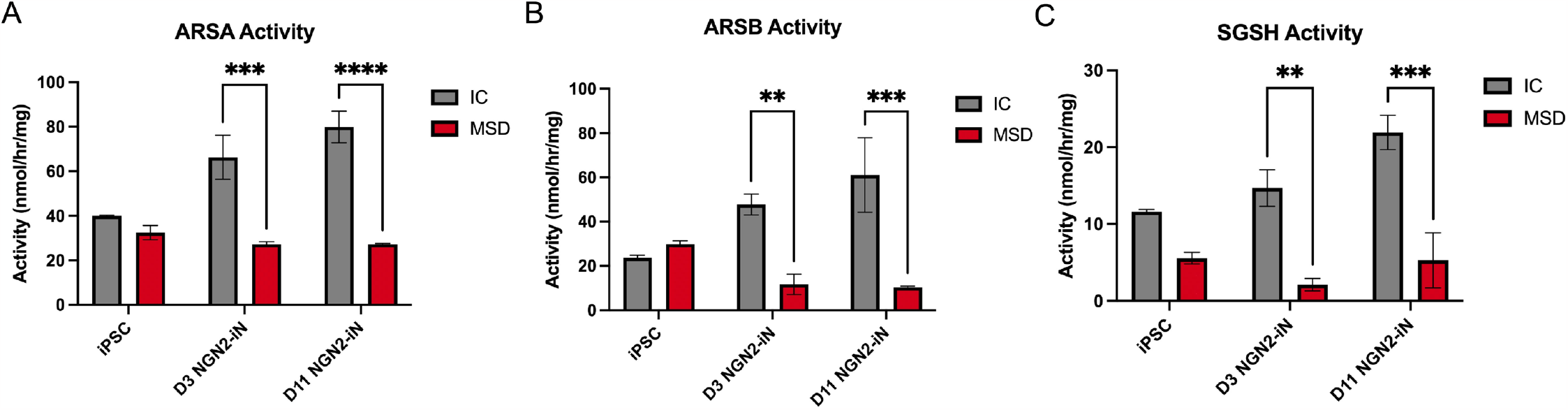
Sulfatase activities in MSD NGN2-iNs are reduced with neuronal maturation. **A**. ARSA, **B**. ARSB, **C**. SGSH activity quantification (nmol/hr/mg total protein) of MSD and isogenic control (IC) cells at different stages of neuronal maturation: iPSCs, Day 3 NGN2-iNs (D3 NGN2-iN), and Day 11 NGN2-iNs (D11 NGN2-iN). Total protein was measured by BCA assay. Data are mean +/-s.e.m. from 3 biological replicates. **, p < 0.01; ***, p < 0.001; and ****, p < 0.0001.

MSD iPSCs did not show significant differences in ARSA and ARSB activities compared to isogenic controls, consistent with the lack of differences seen in *SUMF1* transcript and FGE protein levels (Figs. 5A,B). Interestingly, SGSH activity in MSD iPSCs was significantly decreased compared to isogenic controls (Fig. 5C). At the NPC stage, MSD cells showed trends towards decreased sulfatase activities for ARSA, ARSB, and SGSH (Fig. S3D). In mature neurons, we found that the activities of all three sulfatases were significantly reduced in MSD Day 3 and Day 11 NGN2-iNs compared to isogenic controls, consistent with leukocyte sulfatase activities measured in patients. These results indicate that MSD cells did not experience significant biochemical differences in sulfatase activities until neuronal differentiation is induced. Furthermore, these reduced sulfatase activities may be contributing to the lysosomal stress and maturation defects exhibited by mature MSD neurons.

### 3.6. MSD neuronal cells exhibit GAG accumulation

Functional deficiencies in FGE and downstream sulfatases lead to the accumulation of GAGs in MSD patient cells and animal models. To investigate whether GAGs accumulate in our MSD iPSC, NPC, and NGN-iN cell models, we quantified GAG subspecies by mass spectrometry using the endogenous, non-reducing end (NRE) biomarker method [26,27]. (Figs. 6, S3E). It has been proposed that not all sulfatases are equally vulnerable to loss of sulfatase activity [33]. Previously we found that GAG subspecies associated with mucopolysaccharidoses (MPS) MPS II, IVA, and VI accumulate less as compared to those associated with MPS IIIA and IIID in the urine of MSD patients. Consistent with this, we did not detect significant increases in the GAG species associated with MPS I/II and MPS VI in our MSD cell models (Figs. 6A,B, S3E). MSD NPCs, D3 NGN2-iNs, and D11 NGN2-iNs exhibited minimal concentrations of UA-HexN-UA-2S, a GAG subspecies associated with MPS I/II (Fig. 6A, S3E).

**Figure 6.**
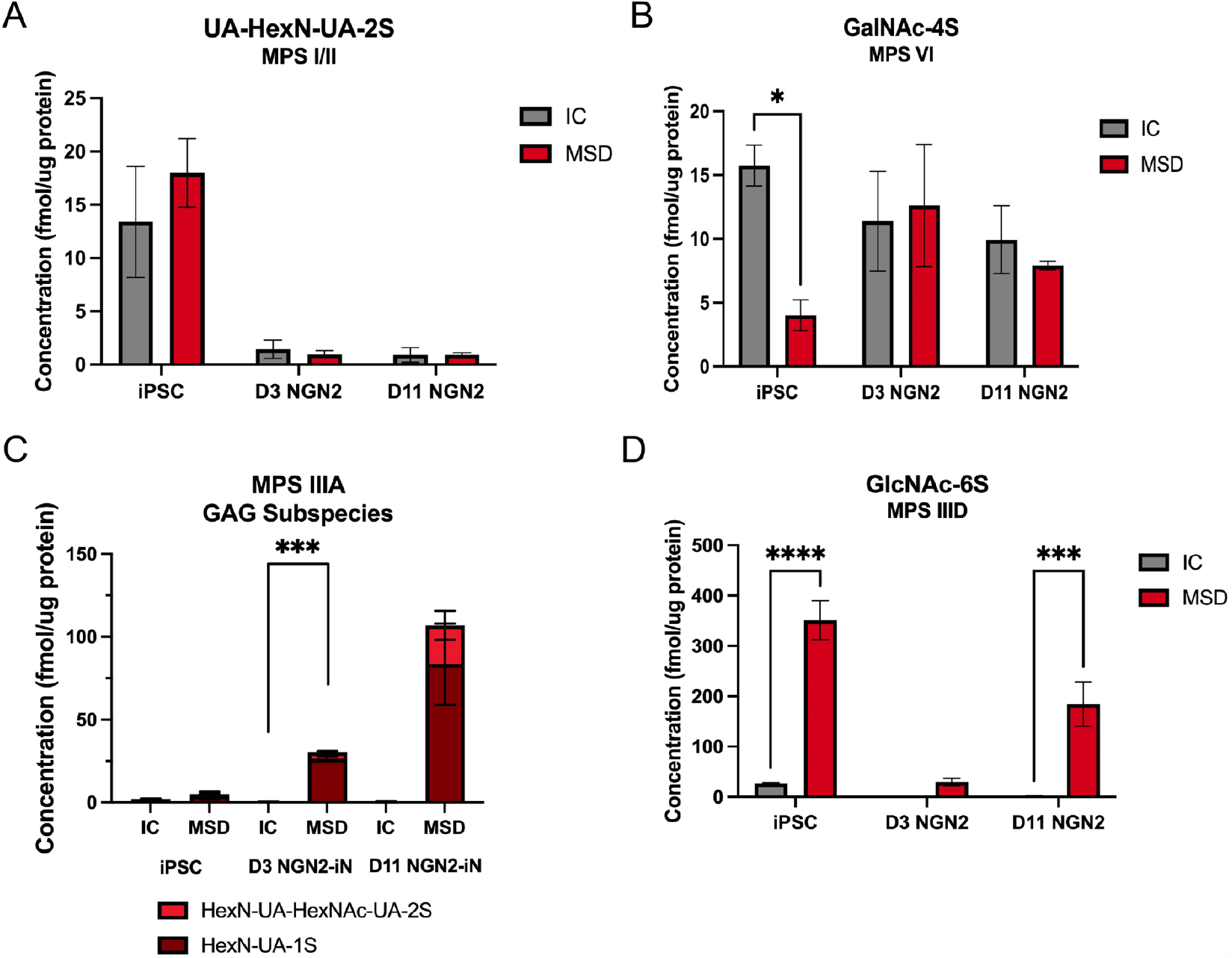
GAGs accumulate in MSD NGN2-iNS with neuronal maturation. **A**. Quantification of GAG subspecies, UA-HexN-UA-2S, associated with MPS I/II as measured by mass spectrometry (endogenous, NRE biomarker method). **B**. Quantification of GAG subspecies, GalNAc-4S, associated with MPS VI. **C**. Quantification of GAG subspecies, HexN-UA-HexNAc-UA-2S and HexN-UA-1S, associated with MPS IIIA. **D**. Quantification of GAG subspecies, GlcNAc-6S, associated with MPS IIID. Concentration of GAGs were normalized to ug of total protein from cell lysates of MSD and isogenic control (IC) cells at different stages of neuronal maturation: iPSCs, Day 3 NGN2-iNs (D3 NGN2-iN), and Day 11 NGN2-iNs (D11 NGN2-iN). Total protein was measured by BCA assay. Data are mean +/-s.e.m. from 3 biological replicates. *p < 0.05, ***p < 0.001, and ****p < 0.0001.

Conversely, MSD patients accumulate high levels of MPS IIIA and IIID GAG subspecies in urine. Our iPSC model recapitulated this accumulation. Specifically, we found that there was a progressive accumulation of GAG subspecies associated with MPS IIIA and IIID with MSD NGN2-iN maturation (Figs. 6C,D). Specifically, HexN-UA-1S and HexN-UA-HexNAc-UA-2S are subspecies associated with MPS IIIA and showed the greatest accumulation in D11 MSD NGN2-iNs compared to isogenic controls (Fig. 6C). There were no differences in these MPS IIIA GAGs between MSD and isogenic controls at the iPSC and NPC stages, as concentrations of these subspecies were all very low (Figs. 6C, S3E). These data suggest that MSD iPSCs and NPCs do not yet accumulate these GAGs, consistent with our cellular and biochemical findings. Of note, we found a significant increase in MSD iPSCs for GlcNAc-6S, a GAG subspecies associated with MPS IIID (Fig. 6D).

Taken together, these findings indicate that when MSD cells are pushed to neuronal maturation there is an accumulation of the GAG species particularly associated with Sanfilippo syndrome (MPS III). This preferential storage accumulation suggests that there may be a sensitivity of specific sulfatases to FGE loss in MSD, especially in neurons.

## 4. Discussion

Generating models that accurately recapitulate human disease is integral to both expand our understanding of disease pathophysiology and develop effective therapies for patients. Here, we generated and characterized a novel human model of MSD by reprogramming patient PBMCs into iPSCs. We also generated an isogenic control iPSC line by correcting the patient mutation using CRISPR/Cas9 gene editing.

Establishing these patient-derived iPSC models enabled us to create two more cellular models of MSD to specifically study disease mechanisms in the brain, a particularly vulnerable organ in MSD. MSD is one of the numerous LSDs with profound CNS involvement in patients. However, the neuropathology of MSD is not well understood. As is the case with many LSDs affecting the CNS, the direct link between lysosomal storage accumulation and neurodegeneration in MSD has yet to be elucidated. Our successful differentiation and characterization of MSD NPCs and NGN2-iNs from iPSCs enabled us to explore the effects of MSD pathology on neuronal cells.

Interestingly, we found a discrepancy between FGE protein levels and *SUMF1* mRNA levels in MSD patient-derived cells. We did not detect significant changes in FGE protein expression in MSD iPSCs, NPCs, or NGN2-iNs. This finding was striking as pathogenic variants of FGE that cause MSD are known to cause the early degradation of FGE [30,31]. However, while FGE protein expression was not significantly decreased compared to isogenic controls, qPCR analysis revealed that *SUMF1* gene expression was significantly reduced in MSD NGN2-iNs at Days 3 and 11 of differentiation. This difference may be a result of a high turnover rate of FGE. For example, FGE variants may still exhibit higher rates of degradation, as suggested by the decreased sulfatase activities exhibited by mature MSD NGN2-iNs. However, increased levels of translation may contribute to compensatory protein levels by Western blot analysis. Further investigation using pulse-chase experiments may be needed to more clearly elucidate the stability of FGE variants in these MSD cell types.

In addition, there was an increasing trend of *SUMF1* mRNA expression with neuronal differentiation in isogenic controls, but not in MSD cells. This finding may suggest an unknown mechanism of mRNA regulation and stability in MSD. *SUMF1* has been found to be tightly regulated by the microRNA, miR-95 [12]. However, because miR-95 is not expressed in mice, its regulation of *SUMF1* has not been well-studied. Our novel human iPSC model can serve as a unique tool to further investigate *SUMF1* mRNA regulation in MSD.

Loss of sulfatase activities in MSD could impair the ability of cells to proliferate or differentiate into mature neurons. At the cellular level, MSD iPSCs and NGN2-iNs showed phenotypic differences in both growth and neurite maturation. Interestingly, MSD iPSCs showed a decreased rate of proliferation as compared to isogenic controls, despite only displaying a functional deficiency in one sulfatase, SGSH. These results may reveal an increased sensitivity of SGSH to FGE loss in iPSCs, leading to MSD phenotypes. Conversely, MSD NGN2-iNs showed deficiencies in all three sulfatase activities measured (ARSA, ARSB, and SGSH). These findings demonstrate that neuronal sulfatases are selectively vulnerable to FGE dysfunction in MSD, as compared to sulfatases in iPSCs and NPCs.

GAG accumulation was significantly increased in MSD NGN2-iNs, specifically for GAG subspecies associated with MPS IIIA and IIID. Notably, we found no significant differences in GAG subspecies associated with MPS I/II and VI between MSD and isogenic control cells in all cell types. These findings are consistent with clinical MSD patient data, where there are no significant differences in the GAG species derived from MPS VI between patients with differing disease severities. On the other hand, there is an elevation in MPS III-derived GAG subspecies in all patients regardless of severity [33].

In conclusion, we generated a patient-derived iPSC model of MSD that is capable of being differentiated into neural cell types to study neuropathology in a human context. The MSD iPSC-derived NGN2-iNs showed significant decreases in *SUMF1* gene expression, impaired neurite outgrowth and maturation, deficiencies in sulfatase activities, and GAG accumulation. This established MSD iPSC model will be an important tool to study pathology in a wide range of cell types because MSD, in addition to being a neurological disorder, affects many organ systems in patients. Furthermore, MSD iPSC-derived cells can be used for future screening and development of potential therapeutics.

## Acknowledgements

This work was supported by a grant from the MSD Action Foundation. We would like to thank Alan Finglas, Amber Olsen, and the MSD patient advocacy community, families, and patients for supporting our work.

## Author Contributions

VP, LSF, MMC, and RAN contributed to the experimental design. VP, LSF, MMC, JAM, ALG, EAW, KK, PW, and XH performed the experiments and collected the data. JAM, ALG, EAW generated, maintained, differentiated iPSCs and iPSC-NPCs. VP, LSF, MMC maintained and differentiated iPSCs and NGN2-iNs. DLF, LS, BLD, and RAN supervised the project. VP and RAN prepared the manuscript with input from LSF, MMC, JAM, ALG, EAW, DLF, XH, LS, and BLD. All authors discussed the results and participated in critical review and revision of the manuscript.

## Ethics Declaration

All procedures followed were in accordance with the ethical standards of the responsible committee on human experimentation (institutional and national) and with the Helsinki Declaration. Informed consent was obtained from all patients included in the study. Individuals with confirmed diagnosis of MSD were enrolled in the Children’s Hospital of Philadelphia IRB-approved protocol #09-007042.

