## Supplemental Figures for "A novel iPSC model reveals selective vulnerability of neurons in Multiple Sulfatase Deficiency"

**Supplemental Figure S1. Characterization of MSD and isogenic control iPSC lines.**
**A.** Karyotyping analysis of MSD iPSC clone revealed normal female karyotype. **B.** Mycoplasma testing of MSD iPSC clone used to generate the iPSC line indicated negative. **C.** Karyotyping analysis of isogenic control iPSC clone revealed normal female karyotype. **D.** Mycoplasma testing of isogenic control iPSC clone used to generate the iPSC line indicated negative. GAPDH was used as a control.

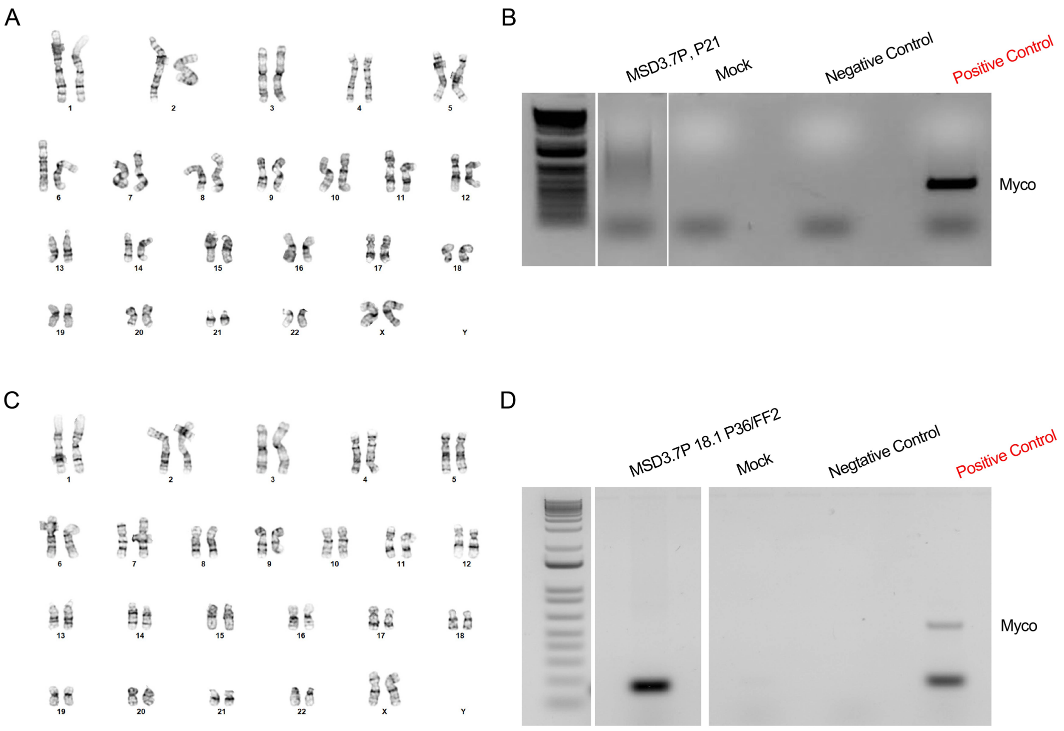

**Supplemental Figure S2. MSD and isogenic control NPCs show characteristic NPC markers.**
**A.** Flow cytometry analysis of MSD NPCs measuring expression of classical stemness markers at Day 0 and NPC markers at Days 8 and 14. **B.** Flow cytometry analysis of isogenic control NPCs measuring expression of classical stemness markers at Day 0 and neuronal maturation markers at Days 8 and 14.

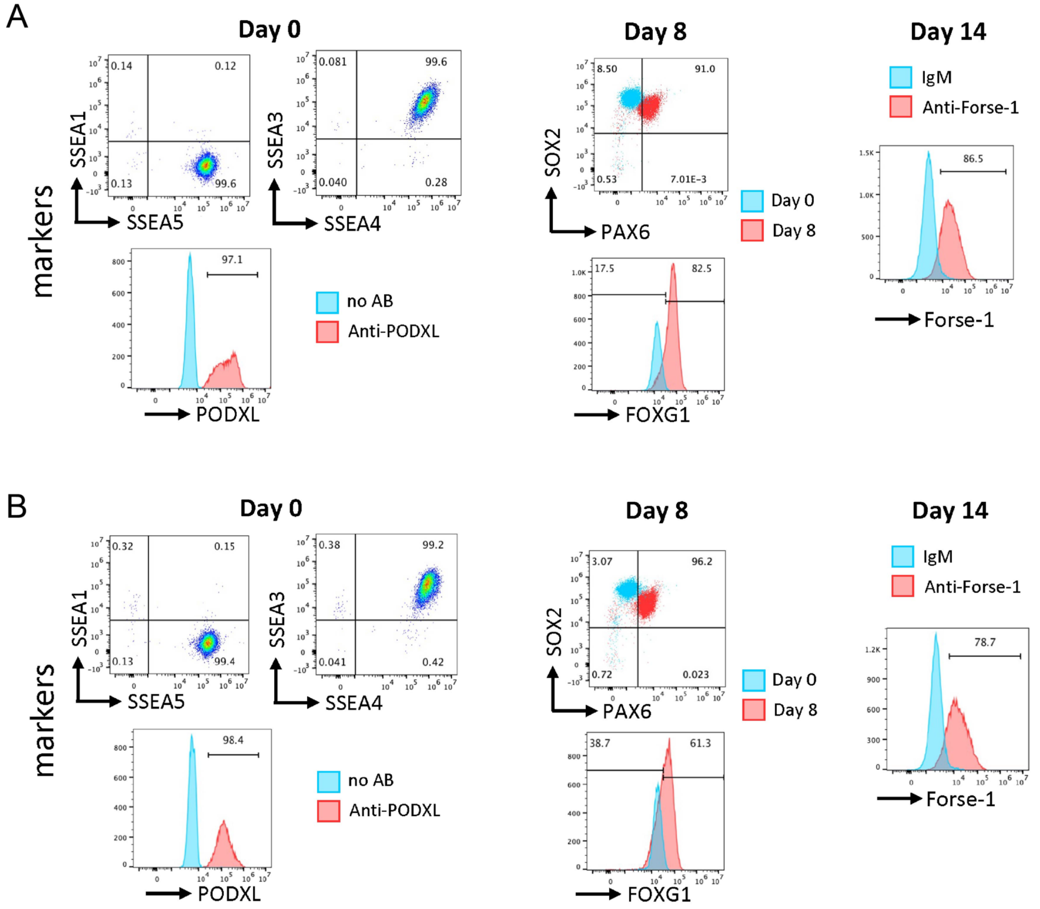

**Supplemental Figure S3. MSD NPCs have reduced sulfatase activities and GAG accumulation.**

**A.** Western blot analysis of FGE protein expression in MSD and isogenic control (IC) NPCs (left). 42 kDa band represents the glycosylated form of FGE and 38 kDa band represents the non-glycosylated form of FGE. Vinculin was used as a loading control. Quantification of FGE Western blot analysis (right). FGE protein expression was quantified as a ratio to the vinculin loading control per sample. Data are mean +/- s.e.m. from 1 biological replicate. **B.** Quantification of *SUMF1* gene expression in MSD and isogenic control NPCs. Relative *SUMF1* mRNA expression was normalized to the average expression of three different housekeeping genes as controls (*GAPDH*, *RNA18SN5*, and *RPLP0*). Data are mean +/- s.e.m. from 3 biological replicates. **C.** Western blot analysis of LAMP1 protein expression in MSD and isogenic control NPCs (left). Vinculin was used as a loading control. Quantification of LAMP1 Western blot analysis (right). LAMP1 protein expression was quantified as a ratio to the vinculin loading control per sample. Data are mean +/- s.e.m. from 1 biological replicate. **D.** ARSA, ARSB, and SGSH activity quantification (nmol/hr/mg total protein) of MSD and isogenic control NPCs. **E.** Quantification of GAG subspecies UA-HexN-UA-2S (MPS I/II), GalNAc-4S (MPS VI), HexN-UA-HexNAc-UA-2S and HexN-UA-1S (MPS IIIA), and GlcNAc-6S (MPS IIID). Concentration of GAGs were normalized to µg of total protein from cell lysates of MSD and isogenic control NPCs.

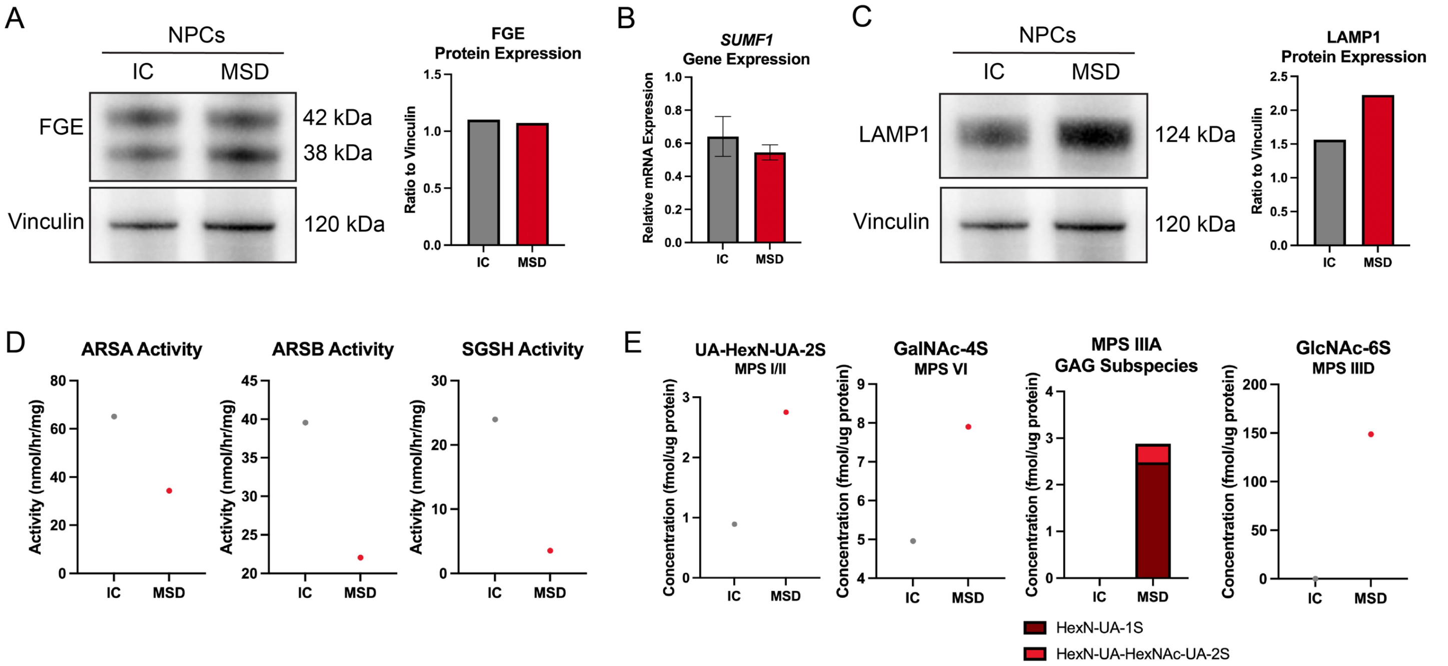
